# Neuronal Activation of the Gαq Protein EGL-30/GNAQ Late in Life Rejuvenates Cognition Across Species

**DOI:** 10.1101/2023.06.06.543909

**Authors:** Morgan E. Stevenson, Gregor Bieri, Rachel Kaletsky, Jonathan St. Ange, Laura Remesal, Karishma J. B. Pratt, Shiyi Zhou, Yifei Weng, Coleen T. Murphy, Saul A. Villeda

**Affiliations:** Lewis-Sigler Institute for Integrative Genomics, Princeton University, Princeton, NJ 08544, USA; Department of Molecular Biology, Princeton University, Princeton, NJ 08544, USA; Department of Anatomy, University of California, San Francisco, San Francisco, CA 94143; Bakar Aging Research Institute, San Francisco, CA 94143

## Abstract

Cognitive decline is perhaps the most devastating aging loss. EGL-30/GNAQ and Gαq signaling pathways are highly conserved between *C. elegans* and mammals. We find that activation of EGL-30 in aged worms at least triples memory span, and we wondered if this highly conserved pathway could also improve memory in very old mice. Murine *Gnaq* is enriched in hippocampal excitatory neurons and declines with age. Furthermore, GNAQ gain-of-function significantly improved memory in aged mice: GNAQ(gf) in hippocampal neurons of 24-month-old mice rescued age-related impairments in health metrics and long-term memory. Single-nucleus RNAseq revealed gene expression changes related to synaptic function, axon guidance, and learning and memory pathways. Several worm orthologs of mouse genes upregulated by GNAQ(gf) overexpression are required for EGL-30(gf)-dependent memory improvement. These results demonstrate that the molecular and genetic pathways between *C. elegans* and mammals are highly conserved, as activation of EGL-30/GNAQ, a pathway first identified in worms, rejuvenates cognitive function in two-year old mice (the equivalent of 70-80 yo humans). To our knowledge, this is the oldest age an intervention has successfully improved age-related cognitive decline.

**One-Sentence Summary:** Neuronal activation of the Gαq protein EGL-30/GNAQ restores long-term memory at old age in worms and mice.

## Introduction

Perhaps the most debilitating feature of aging is the loss of cognitive abilities, especially memory. As in humans, *C. elegans* and mice show learning and memory decline with age (*1*). The cAMP element binding protein (CREB) transcripttion factor is critical for activating the plasticity mechanisms necessary for longterm memory consolidation in vertebrates and invertebrates (*2*–*5*), and CREB expression and function declines with age (*6*).

Several rejuvenation methods, including exercise (*7*–*9*) and parabiosis (*6*) can improve memory in aging animals, and activation of CREB through rejuvenation methods (*6, 10, 11*) or CREB overexpression (*12*) increases synaptic plasticity and improves memory in old age.

EGL-30/G^αq^ is a regulator of pre-synaptic transmission and several functions in *C. elegans* (*13*–*17*), and we found that it also regulates memory (*18*). We previously showed that a constitutively active mutant, *egl-30(js126)* (*19*), significantly extends long-term associative memory (LTAM) lasting more than 24 hours, through the activation of CREB. This enhanced memory was extended by expressing constitutively active forms of EGL-30 (Q205L and V180M) solely in the AWC neuron. Furthermore, *egl-30(gf)* induced in the AWC in mid-life (Day 5) still extended LTAM, when wild-type worms can no longer form longterm memories (*18*). EGL-30 is highly conserved (82% amino acid identity) with mammalian *Gnaq*/GNAQ. Therefore, we wondered if *Gnaq(gf)*, like *egl-30(gf)*, could improve memory and slow cognitive decline in aged mammals, and if so, whether the downstream molecular mechanisms are similarly well conserved across species. Here we found that the analogous activating mutation in GNAQ rejuvenates memory in aged mice through well-conserved neuronal molecular mechanisms.

## Results

### Activated EGL-30 extends memory in *C. elegans* until memory can no longer be measured

We previously found that a gain-of-function mutation (Q205L) in the *C. elegans* Gαq subunit EGL-30 increased memory both in young and aging (Day 5) worms, through the activation of the CREB transcription factor (*18*). However, we did not yet know how late in life EGL-30(gf) induction could rescue memory. By Day 5 of adulthood, wild-type worms have no ability to form long-term associative memories, but can still learn, chemotax, and move (*3*). We used the same heat shock-based induction method to express EGL-30(gf) only in the AWC sensory neurons (*18*) in Day 6, Day 7, and Day 8 worms (**Fig. 1A-C**); we find that in Day 6 and Day 7 worms, 24 hour (LTAM) is fully restored. By Day 8, the worms have difficulty moving, preventing the assessment of memory (**Fig. 1D, E**), since our assay requires the worms to crawl to the butanone spot. That is, while wild-type worms experience the decline of LTAM well before their loss of short-term memory (*3*), induced AWC::*egl-30(gf)* worms retain memory function until they can no longer move well, at least tripling “memory span” (**Fig. 1F**).

**Fig. 1.**
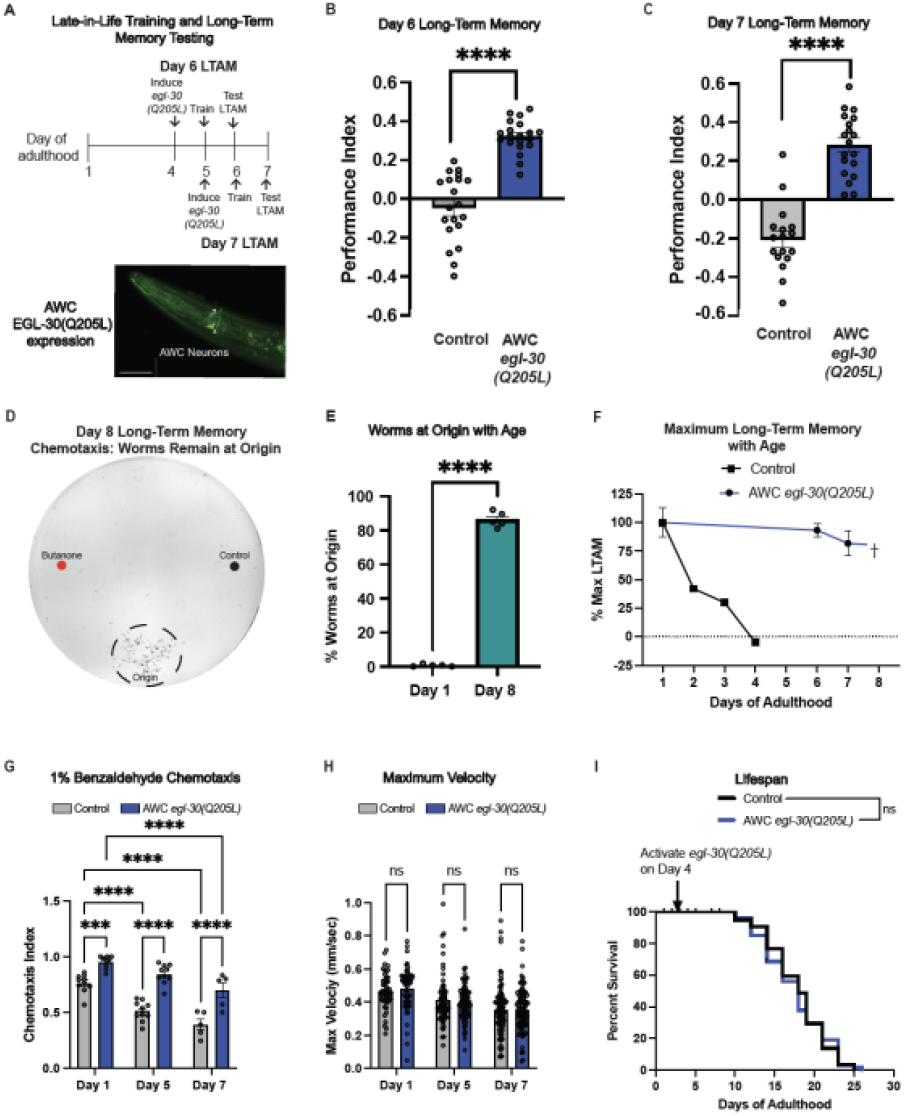
Late-in-life induction of *egl-30(gf)* extends memory in old worms. **(A)** Schematic of experiments. A FLP-inducible GFP-tagged *egl-30(Q205L)* expressed in the AWC neurons was permanently activated after heat shock. To induce *egl-30(Q205L)* late in life, worms were heat shocked, trained for memory, and then tested for long-term associative memory (LTAM) the next day. **(B)** LTAM was extended on Day 6 and **(C)** Day 7. **(D-E)** Representative Day 8 chemotaxis plate. By Day 8, worms lose the ability to move to the odorant spots, instead remain on the origin. **(F)** Late-in-life induction of *egl-30(gf)* in the AWC neuron extends memory until worms can no longer move (Day 7), well beyond wild-type loss of LTAM (Day 4; Kauffman et al., 2010). **(G)** AWC *egl-30(Q205L)* results in a higher preference towards benzaldehyde (AWC-sensed odorant). **(H)** AWC *egl-30(Q205L)* does not affect maximum velocity with age or **(I)** lifespan. For memory assays, each dot represents an individual chemotaxis assay plate. Five technical and three biological replicates were performed for each experiment. For maximum velocity, each dot represents an individual worm. One-way ANOVAs or unpaired Student’s t-tests were performed as appropriate. A Kaplan-Meier analysis with log-rank (Mantel-Cox) method was used to compare lifespans. **** *p* < 0.0001, ns = not significant.

In addition to sensing butanone, the AWC neuron senses benzaldehyde (*20*). Like LTAM, late-in-life expression of activated EGL-30 in the AWC neuron significantly improves chemotaxis toward benzaldehyde with age (**Fig. 1G**) but has no effect on the decline of maximum velocity (**Fig. 1H**) or lifespan (**Fig. 1I**). Therefore, EGL-30(gf) rescues memory – previously the function that is most sensitive to aging - beyond the point when the animals have lost their ability to move.

### *Gnaq* is expressed in the mouse hippocampus and decreases with age

EGL-30 is highly conserved, sharing 82% amino acid identity with mammalian *Gnaq*/GNAQ (**Fig. 2A**). We examined a publicly-available single-cell RNA-sequencing dataset of the adult mouse hippocampus (*21*) and found that *Gnaq* is enriched in excitatory neurons in the dentate gyrus (DG), CA1, and CA3 hippocampal subregions (**Fig. 2B**). We then measured levels of *Gnaq* in young (3 month) and old (24 month) mouse hippocampi (**Fig. 2C**); *Gnaq* levels decrease significantly with age in all hippocampal subfields, including the dentate gyrus, CA1, and CA3 (**Fig. 2C-D**).

**Fig. 2.**
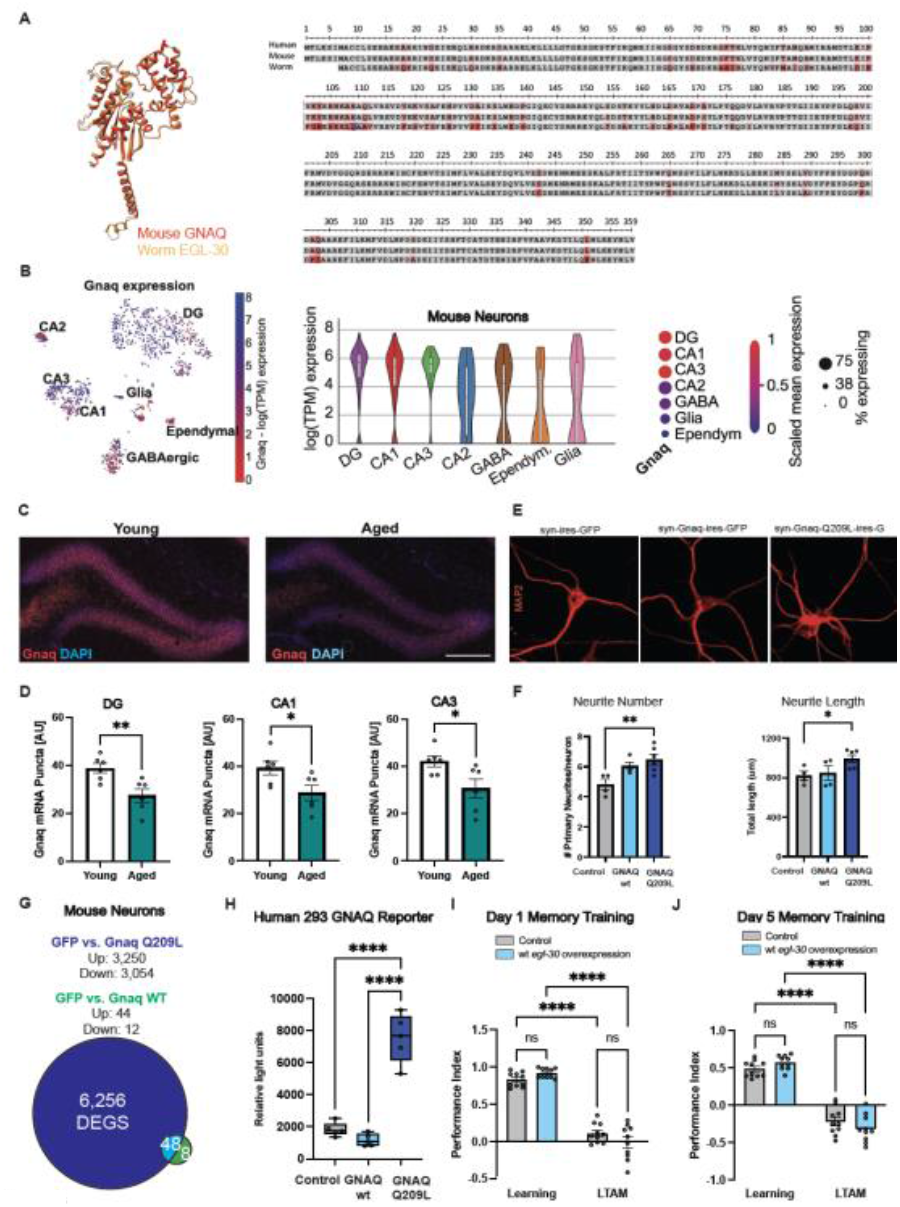
EGL-30 and GNAQ are well conserved, GNAQ declines with age, and the constitutively active mutation is required for extended memory. **(A)** EGL-30 and mammalian GNAQ are highly similar, sharing 82% amino acid identity. Examination of published snRNA sequencing data of the mouse brain (*21*) indicates *Gnaq* is enriched in CA1, CA3, and dentate gyrus (DG). Representative images of *Gnaq* RNAscope in the dentate gyrus. Hippocampal *Gnaq* declines with age when comparing 3-month (young) and 24-month-old (aged) mice. **(D)** Representative images of primary mouse neurons infected with GFP control, wild-type Gnaq, or Gnaq(Q209L). **(E)** GNAQ treatment increased neurite number and length in primary mouse neurons. **(F)** Sequencing of primary neurons transfected with wild-type (wt) GNAQ or GNAQ(Q209L) reveals GNAQ(Q209L) results in more upregulated genes. **(G)** In a human cell line with GNAQ reporter, only GNAQ(Q209L) increases luminescence. **(H)** In worms, wt *egl-30* overexpression does not improve memory in young Day 1 or **(I)** Day 5 aged worms. For mouse experiments, each dot represents an individual mouse. For *in vitro* experiments, each dot represents an individual well. For memory experiments, each dot represents an individual chemotaxis assay plate. One-way ANOVAs or unpaired Student’s t-tests were performed as appropriate. * *p* < 0.05, ** *p* < 0.01, *** *p* < 0.001, **** *p* < 0.0001, ns = not significant.

### Simply overexpressing *Gnaq/egl-30* does not impact neurons or increase memory

Before embarking on tests of *Gnaq* in aged mice *in vivo*, we wanted to determine whether the gain-of-function mutation was necessary to see neuronal effects, or if simply rescuing or overexpressing the wildtype form of *Gnaq/egl-30* is sufficient. The analogous mutation to EGL-30(Q205L) in mammalian *Gnaq*/GNAQ is Q209L, which results in the constitutive activation of *Gnaq* (*22*). We used a cell-type specific viral-mediated overexpression approach in which mature primary mouse neurons were infected with lentiviral constructs encoding either mouse wild-type *Gnaq*, activated *Gnaq*(Q209L) or *GFP* control under the control of the neuron-specific Synapsin1 promoter. While overexpression of wild-type GNAQ in primary mouse neurons did not elicit morphological changes, we found that Gnaq(Q209L) expression increased neuronal complexity, resulting in more primary neurites that are longer in total length (**Fig. 2E-F**). Concordantly, over-expression of wild-type mouse GNAQ in primary mouse neurons induces fewer changes in differential gene expression compared to activated GNAQ(Q209L), which induces a large set of differentially expressed genes (**Fig. 2G; Fig. S1A**). In cells expressing a luminescent reporter for GNAQ pathway activity, wild-type human GNAQ does not activate the pathway to the same degree as expression of the gain-of-function form of human GNAQ(Q209L) (**Fig. 2H**).

Like the mammalian pathway, overexpression of the wild-type EGL-30 in *C. elegans* causes no increase in LTAM on Day 1 or Day 5 of adulthood (**Fig. 2I-J**), unlike expression of the gain-of-function mutation EGL-30(Q205L) (**Fig. 1B-C**). Therefore, the gain-of-function mutation in GNAQ/EGL-30 is necessary both to observe improvements in memory in *C. elegans* and to induce expression changes in the pathway in both mouse and human cells.

### Expression of activated GNAQ increases IEG expression in old mice

To test the possible conservation of EGL-30/GNAQ(gf)’s increased memory function in aged worms in mice, we injected viral vectors expressing either GFP control or GNAQ(Q209L) under the neuron-specific Synapsin1 promoter into the hippocampi of 24-month-old mice (**Fig. 3A-B**). The mice recovered for five weeks, were tested in behavioral assays (see below), and, at ∼26 months old, tissue was collected for molecular analyses (**Fig. 3C**).

**Fig. 3.**
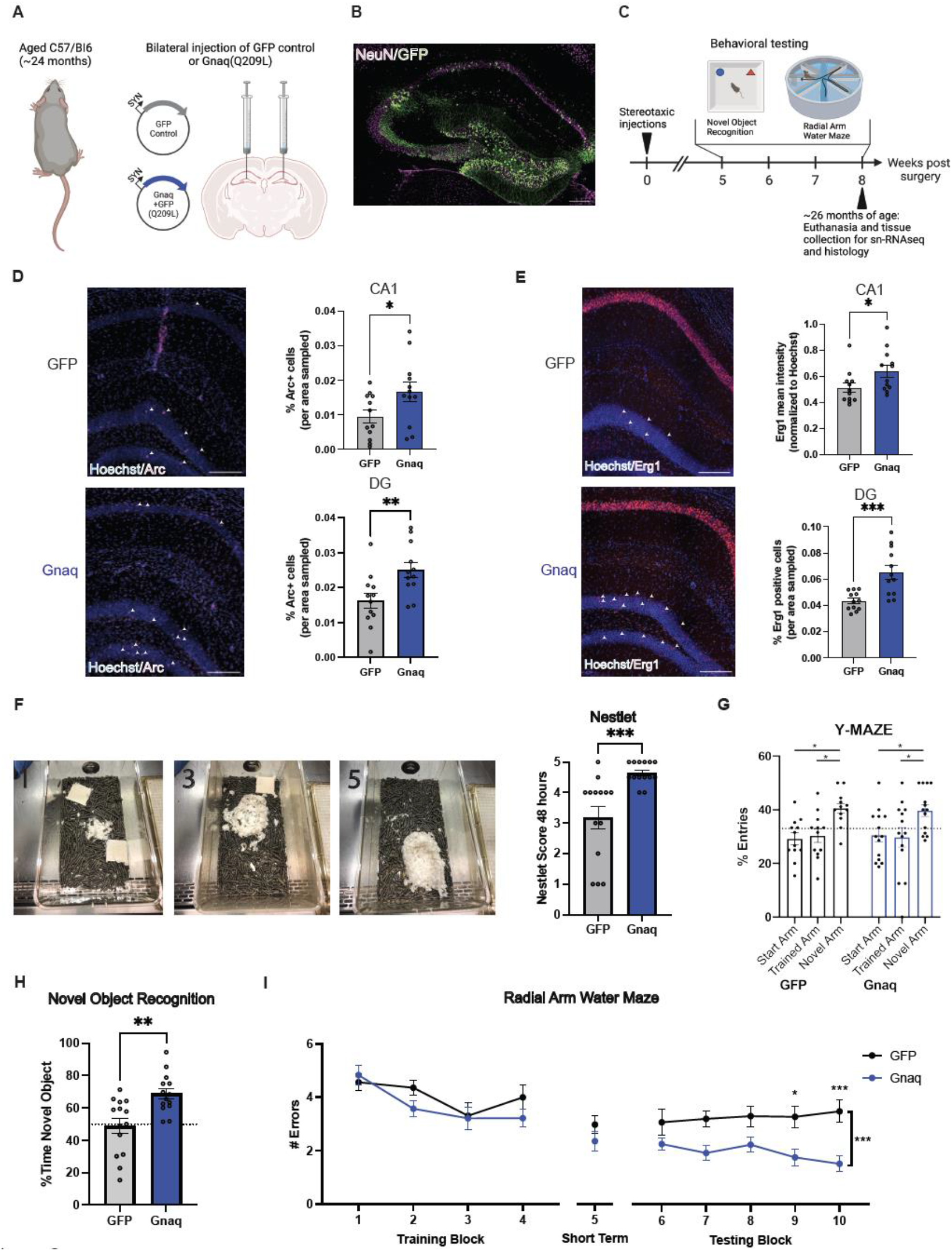
Gnaq(Q209L) increases immediate early gene expression and improves memory in aged mice. **(A)** Schematic of mouse experiments. To test whether *Gnaq(Q209L)* affects behavior, 24-month-old mice were injected with a viral vector leading to the expression of GFP or GNAQ in adult hippocampal neurons **(B). (C)** Aged mice were tested on the novel object recognition and radial arm water maze and euthanized at ∼26 months of age for histology and single nucleus RNA-sequencing. Expression of **(D)** Arc and **(E)** Erg1 were increased in the CA1 and dentate gyrus (DG) regions in GNAQ-treated mice relative to GFP controls (representative immunofluores-cence images). **(F)** Representative nests, where 1 = poor, 3 = average, and 5 = perfect nest. GNAQ-treated mice built significantly better nests than GFP controls, indicating better well-being. **(G)** GNAQ-treated mice showed no differences with GFP control in the Y-maze test (short-term memory). **(H)** GNAQ-treated mice had improved 24-hour long-term memory on the novel object recognition task. **(I)** GNAQ-treated mice had improved long-term memory (blocks 6-10) compared to GFP controls, but there was no difference in learning (training block 1-4) or short-term memory (block 5). Each dot represents an individual mouse. Unpaired Student’s t-tests were performed to compare the two groups. * *p* < 0.05, ** *p* < 0.01, *** *p* < 0.001.

Because *egl-30(gf)* activates CREB and gene transcription activity in *C. elegans* (*18*), we first tested whether there were differences in expression of CREB-regulated immediate early gene (IEG) expression in aged GNAQ(Q209L)-treated mice. Immediate early gene targets *Arc* and *Erg1*, which are downstream of CREB activation and serve as regulators of DNA methylation, neuronal activity, synaptic plasticity, and memory formation (*23*–*25*), are elevated in CA1 and DG in the hippocampi of GNAQ-treated mice (**Fig. 3D-E**).

### Expression of activated GNAQ increases long-term memory in old mice

We next tested the aged mice in behavioral assays. Improved nest-building is considered a metric of better well-being (*26*), and nest building typically declines with age (*27*); GNAQ-treated mice displayed nestlet behavior reminiscent of younger animals (**Fig. 3F**). While there were no differences in short-term working memory, as measured through entries into the novel arm of a Y-maze (**Fig. 3G**), GNAQ(Q209L)-treated mice showed significant improvements in metrics of long-term memory. Specifically, we find that GNAQ-treated mice spend more time with the novel object when tested 24 hours later, indicating better long-term memory (**Fig. 3H**). Perhaps most strikingly, while they showed no differences from controls in the training blocks (learning) or during the short-term memory testing (block 5) of the radial arm water maze (RAWM), the GNAQ-treated mice had significantly better performance in long-term memory (the 6-10 testing block) than did GFP control mice when tested the following day (**Fig. 3I**). That is, like *egl-30(gf)* worms, aged mice with increased expression of *Gnaq(Q209L*) showed improvements in long-term memory. Thus, *Gnaq(Q205L)* expression in old mice has a significant rejuvenating effect, rescuing long-term memory function and increasing quality of life of aged animals.

### GNAQ(Q209L)-treated mice have significant changes in transcription

To determine the cellular and molecular mechanisms that GNAQ(Q209L)-treated mice may use to improve their long-term memory, we performed single-nucleus RNA-sequencing of the hippocampal cells of control and GNAQ-treated mice. All of the major hippocampal cell types were represented in our single-nucleus sequencing data (**Fig. 4A**); activated GNAQ treatment changes gene expression (**Table S1**) but does not cause major changes in cell types (**Fig. 4B**). Differential gene expression analysis showed that the CA1 pyramidal and DG granule neurons – the cells targeted by our intervention – were the sites of the majority of differential gene expression (**Fig. 4B;** examples in **Fig. 4C**). In addition, there were also many upregulated genes in the oligodendrocyte cluster, indicating at least some cell non-autonomous effects of GNAQ treatment (**Table S1**).

**Fig. 4.**
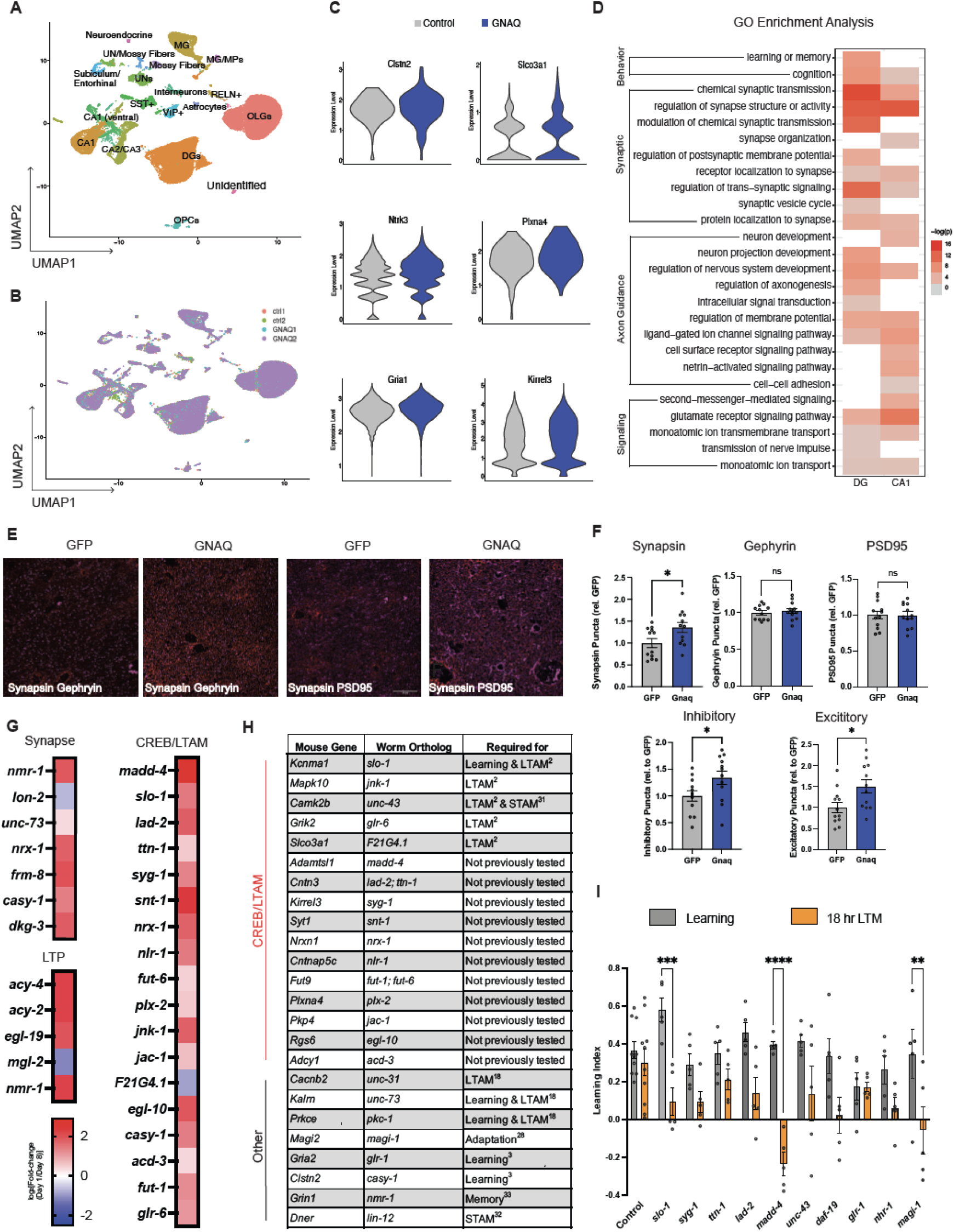
Gnaq treatment upregulates synaptic plasticity genes and increases synapse number and neurite complexity. **(A)** UMAP showing hippocampal cell type clusters. **(B)** No new cell types were induced by GNAQ-treatment. **(C)** Differential expression analysis of significantly upregulated genes in CA1 and dentate gyrus (DG). **(D)** Gene ontology analysis revealed enrichment in CA1 and DG for genes involved in learning and memory, synaptic plasticity, axon guidance, and signaling. **(E)** Representative images of presynaptic (synapsin) and post-synaptic (PSD95, gephyrin) labeling in CA1. **(F)** GNAQ-treatment increases inhibitory and excitatory synapses, with synapsin expression also elevated in CA1. **(G)** Worm orthologs for the GO term category “regulation of synapse structure and activity,” CREB/LTAM genes, and LTP genes are generally higher in young (Day 1) worms, relative to aged (Day 8) worms. **(H)** Many homologs of Gnaq-induced genes identified here overlapped with our CREB-dependent long-term memory list, and are required for worm memory. **(I)** Knocking down worm orthologs (*madd-4, slo-1, magi-1*) in the *egl-30* background with RNAi impairs long-term memory ex7tension. Each dot represents an individual mouse for synapse experiments. Each dot represents one plate for worm experiments. One-way ANOVAs or unpaired Student’s t-tests were performed as appropriate. * *p* < 0.05, ** *p* < 0.01, *** *p* < 0.001, **** *p* < 0.0001, ns = not significant.

### Synapse formation is induced by GNAQ activation

Building on our single-nucleus RNA sequencing analysis indicating an enrichment of synaptic-related changes (**Fig. 4A-D**), we next tested whether GNAQ activation affects synapse number *in vivo* in aged mice. We labeled hippocampal sections from GFP control and GNAQ-treated mice with synapse markers (Synapsin, Gephyrin, and PSD95). Confirming the prediction that increased expression of synapse-structure gene expression might suggest increased synapse formation, we found that activated GNAQ-treated mice have significantly more inhibitory and excitatory synapses than GFP control animals in CA1 (**Fig. 4E-F**), despite the fact that these mice are at least 25 months old. The increase in synapse number appeared to be driven by an increase in presynaptic densities (Synapsin) (**Fig. 4F**). Together, our data suggest that activated GNAQ restores synaptic density, even at an advanced age.

### GNAQ activation upregulates synaptic plasticity, axon guidance, and learning and memory genes

Our differential gene expression analysis revealed an enrichment in synapse organization and structure terms in hippocampal dentate gyrus and CA1 excitatory neurons (**Fig. 4D**). One of the most highly enriched GO term categories was “synapse activity and organization,” which included genes known to be necessary for learning and memory in mammals and *C. elegans*, such as *Clstn2/casy-1, Kalrn/unc-73*, and *Magi2/ magi-1* (*2, 3, 18, 28*–*30*). Our expression analyses in aging worm neurons show that genes from this category decline with age in *C. elegans* (**Fig. 4G**) (Weng & Murphy, unpublished). Other genes known to be required for hippocampal neuron function are also upregulated in the brains of activated GNAQ-treated mice, including genes that function in neurotransmitter /glutamate receptor activity (*Gria1/glr-1, Grik2, Grm1, Grm7/mgl-2*), potassium channels (*Kcnh7/unc-103, Kcnd2/shl-1*), calcium channels (*Cacan1c/egl-19, Cacnb2/ccb-1*), and genes involved in synapse development and function (synaptotagmin *Syt1/snt-1*, neuregulin *Ngr3*, neurexin *Nrxn1/nrx-1, Kalrn/unc-73, Stxbp5/tom-1*, synaptoporin *Synpr/sph-1*, gephyrin *Gphn/moc-1*, and transmembrane proteins such as SynDIG1), cytoskeletal structure and function (*Mid1/madd-2, Carmil1/crml-1, Mast4/kin-4, Arhgef4*), GABA B receptor (*Gabrb1, 2/gab-1*), G protein coupled receptor activity (*Adgrb2*), regulators of CREB activity (Camk2b/*unc-43*, Mapk10/*jnk-1*), solute carrier transporters (*Slc2a13/hmit-1*.*3, Slco3a1 /F21G4*.*1, Slc4a4/ abts-1*) and genes already known to be required for learning and memory (*Tafa2*) (**Fig. 4H**). Surprisingly, genes that play a role early in development, including axon guidance genes (*Dcc/unc-40, Unc5c/unc-5, Slit1, Robo1,2, Dscam, Epha3,6, Sema5a, 6d*) are also upregulated by activated Gnaq.

While many of the GNAQ-upregulated genes are known to be important for neuronal function in mammals, we have also previously found many of the *C. elegans* homologs to be required for long-term associative memory in wild-type worms (e.g., *Slco3a1/F21G4*.*1, Kcnma1/slo-1, Mapk10/jnk-1, Camk2b/unc-43, Kalrn/unc-73, Cacnb2/unc-31*, and *Prkce/pkc-1*) (*2, 18*) (**Fig. 4H)**. Furthermore, *Cacnb2/unc-31*, which is involved in neuropeptide release, is also required for LTAM in the *egl-30* background (*18*). Other genes from our list also overlap with CREB-dependent LTAM genes, including *Adamtsl1/madd-4, Cntn3/ lad-2* and *ttn-1, Kirrel3/syg-1, Syt1/snt-1 Nrxn1/nrx-1, Cntnap5c/nlr-1, Fut9/fut-6, Plxna4/plx-2, Pkp4/jac-1, Rgs6/elg-10, Adcy1/acd-3, Fut9/fut-1*, and *Grik2/glr-6* (**Fig. 4H**)(*2*). Many of these genes also decline with age in worms by Day 8 (**Fig. 4G**), an age when worms no longer have long-term memory (**Fig. 1F**). Others are homologs of genes that are already known to function in *C. elegans* learning or short-term associative memory (e.g., *Magi2/magi-1, Gria2/glr-1, Grin1 glutamate receptor/nmr-1, Dner Notch/lin-12, Camk2b/unc-43*, and *Clstn2/casy-1;* **Fig. 4H**) (*3, 31*–*33*).

### Upregulated GNAQ gene orthologs are required for long-term memory

To test whether the gene expression changes we observed in mice are required for memory, we returned to *C. elegans*, as cognitively testing multiple gene candidates in mice, particularly old mice, would be prohibitively slow. Almost every gene we found to be changed in activated Gnaq-treated hippocampus cells has a well-conserved worm ortholog (**Fig. 4H; Table S1**). In fact, a significant subset of the orthologs were already identified as downstream CREB targets in long-term associative memory (*2*), and of those, we previously found several to be required for memory in wild-type worms (*2*) (**Fig. 4H**).

To test whether activated GNAQ-induced genes (**Fig. 4G-H**) are required for Gnaq/EGL-30 pathway-regulated memory, we used RNAi to knock down specific genes in *egl-30(gf)* adults, trained the animals for food-butanone association on Day 2, and then tested long-term memory 18 hours later (**Fig. 4I)**. We focused on genes involved in synapse activity and organization, declined with age in worms, overlapped with our previously generated CREB/LTAM, or were known to be involved in other forms of memory in *C. elegans* (e.g., *Kcmna/slo-1, Kirrel3/syg-1, Cntn2/ttn-1/lad-2, Adamtsl1/ madd-4, Camk2b/unc-43, Gria2/glr-1, Magi2/magi-1*). We found that several of these genes are required for extended *egl-30(gf)* memory, including *slo-1*/Kcnma1, the alpha subunit of a calcium-activated BK channel; *madd-4*/Adamtsl1, which is involved in axon guidance and synapse organization; and *magi-1*/Magi2, a mem-brane-associated guanylate kinase homo-logue (MAGUK) family member that is involved in habituation and long-term memory (*2, 28*). Thus, we have used *C. elegans* to identify a pathway that rejuvenates memory in aged mice, and the orthologs of gene expression targets altered in the hippocampal cells of these activated GNAQ-treated mice are also required for long-term memory function in the orthologous pathway in *C. elegans*.

## Discussion

Here we have found that a constitutively active form of the murine Gαq protein GNAQ, when expressed in hippocampal neurons of 2-year-old mice, significantly enhances long-term memory. We first identified the role for the EGL-30 pathway in memory in *C. elegans* and here we tested its role in aged mice. These results highlight how well-conserved the molecular and genetic pathways required for memory are from worms to mammals, and suggest that *C. elegans* is a powerful model for the study of memory maintenance with age.

In adulthood, the ability to add new synaptic connections is limited, and this only worsens in old age; however, activated GNAQ treatment rescues these abilities. Differential gene expression and gene ontology analyses suggest that GNAQ(gf) treatment rejuvenates memory function of aged mice through the coordinated expression of genes that increase synaptic structure and function, axon guidance, and neuronal activity. GNAQ intervention targeted adult neurons in the aged hippocampus, and improving function without requiring any obvious markers of stem cell activation, as there is no evidence of neurogenesis in GNAQ-treated mice, suggesting that the memory improvements we observed are the result of changes to existing neurons. Furthermore, there we find no activation of microglia or neuroinflammatory responses in GNAQ-treated mice, suggesting that the treatment is not toxic to the animals. Excitingly, the degree of memory improvement is on the scale of that seen in aged mice with parabiosis and exercise interventions (*6, 9*), which may function in parallel to the GNAQ pathway.

GNAQ treatment upregulates genes that play central roles in consolidation of memory. Single-nucleus RNA sequencing and subsequent gene ontology analysis revealed an enrichment of genes involved in synapse organization and activity, including long-term potentiation. For example, several genes that are directly involved in synaptogenesis and spineogenesis, such as *Nrxn1/nrx-1*, were upregulated in activated GNAQ-treated mice. *Nrxn1/nrx-1* is a critical regulator of synaptic function and synaptogenesis in mice (*34, 35*) and *C. elegans* (*36*). Recently, mouse dentate gyrus neurons involved in encoding memory were found to have a late-onset increase in *Nrxn1* mRNA splicing; activity-dependent histone modifications regulated this *Nrxn1* splicing, which was required for preserving contextual fear memories (*37*). CAMKIIB/ UNC-43 also interacts directly with the cytoskeleton, and regulates spineogenesis and maintenance in adulthood via its interactions with F-actin in rodents (*38*), and is important for short-term and long-term memory in *C. elegans* (*2, 31*). CAMKIIB’s structural role at the synapse is important for Schaffer collateral-CA1 synaptic plasticity and hippocampal learning (*39*). We also found genes that regulate synapse activity, many of which have direct involvement in long-term potentiation, were upregulated; for example, *Grin1/nmr-1* encodes for GluN1, an essential component of the NMDA receptor, and alternative splicing of *Grin1* exon 5 controls the magnitude of long-term potentiation in CA1, and thus the ability to learn and remember (*40*). *Gria1* encodes for the GluA1 AMPA receptor subunit, and reduction of GluA1 in CA2/CA3 impairs short-term memory (*41*).

Axon guidance genes, including Semaphorins and multiple Netrin, Ephrin, and Slit receptors, are also upregulated in activated GNAQ-treated mice. Although axon guidance genes have been most examined in the context of development, they can guide synaptic plasticity in adulthood. Netrin-1 infusion can recover long-term potentiation and memory impairments that result from amyloid-beta treatment in mice (*42*), and ephrin receptors are involved in long-term potentiation, promote neurotransmitter release, and regulate spine morphology and synapse formation (*43, 44*). However, the roles of these proteins in aged animals have not been previously probed. The induced expression of these axon guidance genes following GNAQ treatment supports our findings of improved memory and synapse density.

It is interesting to consider whether activated GNAQ treatment in neurons could also improve oligodendrocyte function. Although activated GNAQ treatment primarily affected neurons, we also observed an enrichment in genes involved in signal transduction and synaptic transmission in oligodendrocytes. Myelination is inhibited in aged mice, and loss of myelin impairs spatial memory in young mice (*45*). Activation of existing oligodendrocytes can also enhance myelin thickness and conduction speed, leading to better hippocampal memory (*46*).

Testing in aged animals allows the discovery of beneficial mechanisms and acute treatments that might be deleterious with chronic administration or in younger animals. The pathways that *C. elegans* neurons use to function are highly conserved with molecular pathways in mouse neurons, which allowed us to use worms for rapid discovery and testing of possible mechanisms that can rejuvenate aged mammalian neurons. Distinct from the negative effects induced by chronic Gq activation of ventral hippocampal networks early in life (*47*), late-life, temporally-controlled acute activation of Gαq/GNAQ may offer a mechanism to slow or even reverse memory decline in aged individuals.

To our knowledge, GNAQ treatment in two-year old mice is the latest age at which a rejuvenation method has improved memory. Mice between 18-24 months are considered old, but cognition gets still worse in older animals. For example, 22-month-old mice were more impaired on the Morris water maze than 18-month-old mice (*48*). Here, we treated mice with activated GNAQ when they were 24 months old, and they were tested for learning and memory ability when they were 25-26 months old. This is a remarkably old age to employ rejuvenation methods and test cognition in mice, as most previous experiments begin interventions and cognitive testing between 18-20 months old, including treatments that continue during memory training. For example, exercise (*7*) and treatment with parabiosis or plasma factors from young or exercised mice (*6, 9*) improve memory when interventions begin around 18 months of age. The effects of young cerebral spinal fluid on aged mouse memory was tested around 20 months of age (*49*). Similarly, transplantation of young microbiota improved longterm memory in 19-20 month old mice (*50*). One month of environmental enrichment improves memory in mice as old as 21 months of age (*51*). Treatment with ISRIB, a drug-like small-molecule that inhibits the integrated stress response, improves memory in 19 month old mice (*52*), but later ages were not tested. Therefore, the fact that activated GNAQ treatment significantly improves cognition in mice more than two years old is unprecedented, as far as we know.

Activated GNAQ treatment upregulates synaptic plasticity mechanisms that improve memory across organisms. Moreover, these activated GNAQ-treated mice display signs of improved well-being and increased synapse formation. GNAQ intervention late in life can rejuvenate post-mitotic neurons in the aged hippocampus, producing an environment that supports plasticity and in turn memory consolidation, with no evidence of neuroinflammation or microglia activation. The fact that memory in such aged mice – the equivalent to 70–80-yearold humans – could be rescued by activated GNAQ treatment suggests that aged neurons may have more potential to remodel than has been previously recognized.

## Acknowledgments

We thank the Princeton FACS Core and Christina DeCoste for her assistance, the Princeton Confocal Microscopy Core, the Princeton Genomics Core and Jennifer Miller and Jean Arly Vomar for their assistance, the C. elegans Genetics Center, and members of the Murphy lab for suggestions on the manuscript. We acknow-ledge the UCSF Parnassus Flow Core (RRID:SCR_018206), UCSF Center for Advanced Technology, and the Stanford Sherlock cluster. We also thank the Murphy and Villeda labs for feedback on this manuscript.

## Funding

Funding for this work was provided by the Simons Collaboration on Plasticity in the Aging Brain to CTM and to SAV; Lev Mikheev, an NIH F32 award to MS (AG079490) and GB (AG081038); a Hillblom Postdoctoral Fellowship to GB, NSF Pre-doctoral award to JSA, an NIA R01 (AG077816) to SAV, and an NIH Director’s Pioneer Award to CTM (DP1AG077430).

## Author contributions

Conceptualization: CTM, SAV

Methodology: MES, GB, RK, JSA, KJBP, LR, CTM, SAV

Investigation: MES, GB, RK, JSA, KJBP, LR, SZ, YW

Visualization: MES, GB, RK, JSA, LR, YW

Funding acquisition: CTM, SAV

Project administration: CTM, SAV

Supervision: CTM, SAV

Writing – original draft: MES, RK

Writing – review & editing: CTM, SAV

## Competing interests

CTM and SAV have no competing interests.

## Data and materials availability

With the exception of single-nucleus RNA-sequencing and mouse neuron sequencing data, all other data are available in the main text or the supplementary materials. Sequencing data are available at BioProject #PRJNA972702.

### Methods

### *C. elegans* Experiments

#### *C. elegans* maintenance

Strains were maintained at 20°C for the duration of experiments. Maintenance plates were made with high growth medium (HGM: 3 g/L NaCl, 20 g/L Bacto-peptone, 30 g/L Bacto-agar in distilled water, with 4 mL/L cholesterol (5 mg/mL in ethanol), 1 mL/L 1M CaCl_2_, 1 mL/L 1M MgSO_4_, and 25 mL/L 1M potassium phosphate buffer (pH 6.0) added to molten agar after autoclaving). All assays were performed on plates made with standard nematode growth medium (NGM: 3 g/L NaCl, 2.5 g/L Bacto-peptone, 17 g/L Bacto-agar in distilled water, with 1 mL/L cholesterol (5 mg/mL in ethanol), 1 mL/L 1M CaCl_2_, 1 mL/L 1M MgSO_4_, and 25 mL/L 1M potassium phosphate buffer (pH 6.0) added to molten agar after autoclaving (*53*). All experiments that did not involve RNAi treatment were seeded with OP50 *E. Coli* (BactoBeads) for *ad libitium* feeding. Hypochlorite synchronization was used to develop-mentally synchronize experimental worms, where gravid hermaphrodites were exposed to an alkaline-bleach solution (e.g. 6 mL sodium hypochlorite, 2.5 mL KOH, 41.5 mL distilled water) to collect eggs, followed by repeated washes with M9 buffer (6 g/L Na_2_HPO_4_, 3 g/L KH_2_PO_4_, 5 g/L NaCl and 1 mL/L 1M MgSO_4_ in distilled water (*53*)

#### FUdR treatment

For aging experiments, worms were transferred at the L4 larval stage to HGM plates supplemented with 500μl/L 0.1M FUdR (5-Fluoro-2’-deoxyuridine; Sigma Aldrich, Cat. # F0503) for a final concentration of 0.05M FUdR at the L4 larval stage. Worms were moved off FUdR plates, back to HGM plates, 24 hours prior to behavioral assays.

#### RNAi treatment

For adult only-RNAi experiments, worms were transferred at the L4 larval stage to HGM plates supplemented with 1 mL/L 1M IPTG (isopropyl β-d-1-thiogalacto-pyranoside) and 1 mL/L 100 mg/mL carbenicillin. Plates were seeded with HT115 *E. coli* for *ad libitium* feeding and induced with 200 μl of 0.1 M IPTG. RNAi treatment was performed using standard RNAi feeding methods. Worms were maintained on HGM RNAi plates until Day 2 of adulthood. All bacterial clones, including the control vector (pL4440), were sequenced prior to use.

#### Strains

*C. elegans* strain N2 var. Bristol: wild-type/CGC/N2

*C. elegans* strain NM1380:*elg-30(js126)*/CGC/NM1380

*C. elegans* strain CQ429: *pHSP16-48::FLPase;Podr-1::FRT::egl-30(Q205L);*

*Pmyo-2::mCherry/* Arey et al., 2018/CQ429

*C. elegans* strain: CQ701: wtEx74 Topo-Prab-3::GFP-egl-30(WT)::rab3 UTR; *Pmyo-2::mCherry*/this paper/CQ701

*C. elegans* strain CQ601:*egl-30(Q205L);vls69[pCFJ90(Pmyo-*

*2::mCherry+Punc-119::sid-1)]/*this paper/CQ601

#### Construction of Transgenic Lines

Extrachromosomal transgenic arrays were generated for experiments testing the effects of wild-type EGL-30 overexpression. Line CQ701: wtEx74 *Prab-3*::*GFP-egl-30*(WT)::*rab-3* UTR; *Pmyo-2::mCherry* was generated by injecting wild-type worms with 25 ng/μl of *Prab-3*::*GFP-egl-30*(WT)::*rab-3* UTR and 1 ng/μl *Pmyo-2::mcherry. Prab-3*::*GFP-egl-30*(WT)::*rab-3* UTR was made by Gibson cloning of 1215bp of the *rab-3* promoter upstream of *gfp* N-terminally fused to the *egl-30* cDNA, followed by the *rab-3* 3’UTR into a Topo backbone.

#### Heat-shock inducible EGL-30 strain

For experiments using CQ429: *pHSP16-48::FLPase;Podr-1::FRT::egl-30(Q205L); Pmyo-2::mCherry* (*18*), transgenic and non-transgenic siblings were heat-shocked at 34°C for 1 hour to permanently activate the FLP-inducible GFP-tagged gain-of-function EGL-30 in the AWC of transgenic worms (Fig. S1). The heat-shock always occurred 24 hours prior to behavioral assays, to allow worms to recover. The heat-shocked, non-transgenic siblings served as the controls for all experiments. To verify GFP-tagged gain-of-function EGL-30 was expressed in the AWC, a subset of worms were imaged using a Nikon AXR confocal microscope with a 60x oil objective (**Fig. 1A**).

#### Recombinant DNA

Plasmid: pL4440 RNAi control/Ahringer Plasmid: pL4440-*slo-1* RNAi/Ahringer Plasmid: pL4440-*syg-1* RNAi/Ahringer Plasmid: pL4440-*ttn-1* RNAi/Ahringer Plasmid: pL4440-*lad-2* RNAi/Ahringer Plasmid: pL4440-*madd-4* RNAi/Ahringer Plasmid: pL4440-*unc-43* RNAi/Ahringer Plasmid: pL4440-*glr-1* RNAi/Ahringer Plasmid: pL4440-*magi-1* RNAi/Ahringer

#### Positive olfactory associative memory assay

Worms completed short-term associative memory assays and were trained and tested as described previously (*2, 3, 18*). For the short-term memory training, when worms were at the appropriate age of adulthood, they were washed from HGM or HGM+RNAi plates with M9 buffer, allowed to settle by gravity and washed a total of 3x. Next worms completed a 1 hour starvation in M9 buffer followed by one, 1 hour CS-US pairing, where worms were transferred to NGM plates seeded with OP50 *E. Coli* with 18 μl of 10% 2-butanone (Acros Organics; Cat. # 332828-25ML) in ethanol streaked on the lid at 20°C. After conditioning, trained worms were tested for chemotaxis to 10% butanone vs. ethanol control either immediately after training (0 hr/learning) or held on seeded NGM plates at 20°C with fresh OP50 *E. Coli* for 1-24 hours. Short-term memory (1 hour) and long-term memory (18 or 24 hours) were tested by chemotaxis to 10% butanone vs. ethanol control. This is compared with the naïve chemotaxis index to butanone, completed on a subset of worms that did not complete the memory training. Standard chemotaxis assay conditions were used (*20*). If fewer than 20 worms left origin, the plate was excluded from analysis All memory assay experiments included five technical replicates and three biological replicates..

For each plate, chemotaxis indices were calculated as follows: (#worms_Butanone_ – # worms_Ethanol_)/(#worms_total_ – #worms_origin_). Performance indices were calculated to account for naïve butanone chemotaxis: Chemotaxis Index_Trained_ – Chemotaxis Index_Naïve_. For extrachromosomal transgenic strains, chemotaxis indices were manually counted for fluorescent transgenic and non-fluorescent wild-type siblings at individual timepoints on the chemotaxis plates.

#### Chemotaxis assay

Chemotaxis towards 1% benzaldehyde (Millipore Sigma, Cat. # B1334-100G) in ethanol was accessed with age in synchronized Day 1, Day 5, and Day 7 *pHSP16-48::FLPase;Podr-1::FRT::egl-30(Q205L); Pmyo-2::mCherry* transgenic worms or their wild-type siblings using standard chemotaxis assay conditions (*20*). Assays included five technical replicates and three biological replicates.

#### Maximum velocity measurement

Maximum velocity was assessed for *pHSP16-48::FLPase;Podr-1::FRT::egl-30(Q205L); Pmyo-2::mCherry* transgenic worms or their wild-type siblings with age (Day 1, Day 5, Day 7). Worms were picked off HGM plates onto unseeded 60 mm NGM plates for the analysis, avoiding the carryover of any bacteria to the assay plate. Movements were recorded for 30 sec at a rate of 25 frames per second using a Nikon DS-Fi3 camera mounted on a Nikon Smz645 Stero microscope and recorded with Nikon Elements software. Locomotion velocity was recorded as mm/sec. Recorded images were analyzed by ImageJ and wrMTrck (plugin for ImageJ). Only worms that were in the field of view for at least 15 sec were included in the analysis. Data was imported into an Excel spreadsheet and the peak locomotion velocity observed was used for maximum velocity. *n* > 47 worms per day/group.

#### Lifespan analysis

Lifespan analyses were completed for *pHSP16-48::FLPase;Podr-1::FRT::egl-30(Q205L); Pmyo-2::mCherry* transgenic worms or their wild-type siblings to test of late-in-life *egl-30* gain-of-function only in the AWC affected lifespan. At the L4 larval stage worms picked for the assay. Every other day, worms were transferred to freshly seeded NGM plates. The first day of adulthood was defined at T=0. To permanently activate *egl-30* in the AWC late-in-life, all worms were heat-shocked at 34°C for 1 hour on Day 4 of adulthood. Kaplin-Meier analysis with log-rank (Mantel-Cox) method was used to compare lifespans between transgenic and non-transgenic siblings. Worms that “escaped” or “bagged” were censored on the day of the event. *n =* 96 per strain, three biological replicates were performed.

#### Human cell line GNAQ luciferase report assay

The GloResponse NFAT-RE-*luc2P* HEK293 Cell Line (Promega, Cat. # E8510) was used to compare luciferase activity between control GFP, GNAQ wild-type, and GNAQ(Q209L) expressing cells. Cells were grown in at 37°C with 5% CO_2_ in DMEM, high glucose medium (Gibco 11965118; Thermo Fisher Cat. # 11-965-118) with 10% FBS (Thermo Fisher, Cat. # 16140089) and Penicillin-Streptomycin-Glutamine (Thermo Fisher Cat. # 10378016). Cells were plated in white-bottom, 96-well plates (Falcon 353296, Fisher Scientific Cat. # 09-771-26) at a concentration of 2×10^4^ and transfected the following day with the pcDNA3.1-mGreenLantern control (Addgene #161912), pcDNA3.1-human GNAQ WT, and pcDNA-3.1-human GNAQ (Q209L) following the Lipofectamine 3000 Transfection Reagent (Thermo Fisher, Cat. # L3000008) protocol. Forty-eight hours later, the luciferase activity was determined using the Bright-Glo Luciferase Assay System (Promega, Cat. # E2620) reagents, which were prepared following manufacturer instructions and added 8 min prior to reading the plate on a Biotek Synergy Mx plate reader. Luminescence (relative light units) was recorded and averaged within conditions, and three biological replicates were completed.

### Mouse Experiments

#### Animal models

The C57BL/6 mouse line was used for all experiments (The Jackson Laboratory and National Institutes of Aging). All other studies were performed with male mice. The numbers of mice used to result in statistically significant differences was calculated using standard power calculations with α = 0.05 and a power of 0.8. We used an online tool (http://www.stat.uiowa.edu/~rlenth/Power/index.html) to calculate power and sample size based on experience with the respective tests, variability of the assays, and inter-individual differences within groups. Animals utilized for each individual experiment were from the same vendor and aged together. All animals from Jackson Laboratories were acquired at two months of age and aged in-house. Animals were moved to a new location for behavioral assessment at the Villeda Lab Behavioral Suite. Mice were housed under specific pathogen-free conditions under a 12-hour light-dark cycle, and all animal handling and use was in accordance with institutional guidelines approved by the University of California San Francisco IACUC.

#### RNAscope for *Gnaq* expression in young and aged mice

RNAscope was used to compare *Gnaq* expression in young (3 mo. old) and aged (24 mo. old) mice. *In situ* hybridization experiments and fluorescent staining for Gnaq was performed using the RNAScope Multiplex Fluorescent Reagent Kit V2 and the RNA-Protein Co-Detection Ancillary Kit (ACD Bio, Cat. # 323100 and 323180). RNAscope processing followed the manufacturer’s instructions. Cryo-preserved 40 μm thick coronal sections that included the hippocampus were washed in PBS and pre-treated in 200 μl of hydrogen peroxide provided by the RNAscope reagent kit. Sections were treated for 10 min with RNAScope Protease III, incubated for 2 hours with Gnaq RNAScope probe (Cat. # 1067091-C1) in a HybEZ oven set at 40°C. After amplification steps, mouse Gnaq transcripts were detected using TSA Plus Cy3 reagents (Akoya Biosciences, Cat. # SKU NEL744001KT). Sections were placed on slides, coverslipped with Prolong Diamond (Molecular Prgoes, Thermo Fisher, Cat. # P35970) and stored at 4°C. Slides were imaged within 2 weeks using a Zeiss LSM 800 confocal microscope with 20x objective. For quantifications, stacks of 1 μm thick were acquired from the dentate gyrus, CA1, and CA3 subfields of the hippocampus. The number of puncta per ROI were counted as previously describe (*54*) and averaged from 3 hippocampal sections per animal; *n = 6* mice/group.

#### Primary neuron cultures (for sequencing and neurite analysis)

E17 mouse embryo (C57B1/6J) brains were used to dissociate primary mouse hippocampal neuron suspensions using the papain dissociation system (Worthington, Cat. # LK003153). Neurons were seeded on poly-L-lysine (Sigma Aldrich, Cat. # P6282) coated plates [0.1% (wt/vol)] and grown in a humidified chamber at 37°C with 5% CO_2_. Neurobasal medium (Thermo Fisher, Cat. # 21103049) that was supplemented with B-27 serum free supplement (Thermo Fisher, Cat. # 17504044), GlutaMAX (Thermo Fisher, Cat. # 35050061) and penicillin-streptomycin (Thermo Fisher, Cat. # 15140122). Half-media changes were completed every 4-5 days. Neurons were plated on 12 mm glass coverslips (Carolina Biological Supplies Cat. # 633009) in 24-well plates (100,000 cells/well). For immunocytochemistry, cells were fixed for 10 min with 4% paraformaldehyde. Cells were then washed and stained with MAP2 antibody (Sigma-Aldrich, Cat. # M1406) and AlexaFluor 555 conjugated secondary antibody (Thermo Fisher Cat. # A31570). For the neurite analysis, 5 randomly selected image stacks were acquired (10 slices spaced 1.5 μm apart) for each coverslip using a Zeiss LSM900 confocal microscope with a 20x objective. Neurite length and number was assessed using the NeuronJ ImageJ plug. 20-25 neurons were traced and averaged for each coverslip.

#### Bulk RNA-sequencing analysis

Total RNA was isolated from primary neuron cultures using Trizol extraction combined with columns from PureLink RNA Mini kit (ThermoFisher, Cat# 12183025) following the manufacturer’s instructions. To quantify mRNA expression levels, equal amounts of cDNA were synthesized using the HighCapacity cDNA Reverse Transcription kit (ThermoFisher, Cat# 4374966) and mixed with the KAPA SYBR Fast mix (Roche, Cat# KK4601), and Gnaq-specific primers (GCCTACACAACAAGACGTGC, GACCTTTGGCCCCCTACATC). GAPDH mRNA was amplified as an internal control. Quantitative RT-PCR was carried out in a CFX384 Real Time System (Bio-Rad). RNA-sequencing libraries were generated using and sequenced using Genewiz RNA-seq services (New Jersey). Alignment of RNA-sequencing reads to the mouse mm10 transcriptome was performed using STAR v2.7.3a (*55*) following ENCODE standard options, read counts were generated using RSEM v1.3.1(*56*), and differential expression analysis was performed in R v4.0.2 using the DESeq2 v1.28.1(*57*). Detailed pipeline v2.1.2 and options available on https://github.com/emc2cube/Bioinformatics/.

#### Viral vectors

The lentiviral murine Gnaq overexpression construct was generated in a two-step cloning process. First, the Gnaq coding sequence (CDS) and part of the 5’ and 3’ untranslated regions (UTRs) was amplified from adult mouse hippocampal cDNA and cloned into the pENTR-D-TOPO vector (Thermo Fischer, cat# K240020). After sequence validation, the CDS was further PCR amplified and restriction enzyme sites (NheI and BamHI) were incorporated into the forward and reverse primers. Gnaq was then ligated into a Synapsin promoter-based IRES eGFP lentiviral plasmid using traditional restriction enzyme-based cloning strategy. The Q209L mutation was generated using the QuikChange Lighting site-directed mutagenesis kit (Agilent). A Synapsin-GFP construct based on the same lentiviral plasmid was used as a control(*58*). All plasmids were validated by Sanger sequencing prior to virus production. Lentiviral particles were generated as previously described(*59*). Briefly, HEK293T cells were lipotransfected with 4:3:1 mg of lentiviral vector: psPax2:pCMV-VSVG: (psPAX2 was a gift from Didier Trono (Addgene plasmid # 12260); pCMV-VSV-G was a gift from Bob Weinberg (Addgene plasmid # 8454, RRID:Addgene_8454). After 48 hours lentivirus-containing media was centrifuged for 5 min at 1000g and the supernatant filtered through a 45μm filter to remove cellular debris. Media underwent utracentrifugation (24000 RPM for 1.5 hours) to concentrate virus. Viral pellets were gently resuspended in PBS. Lentiviral titers were between 1-3 × 10^9 viral particles per mL. Primary neuron cell cultures were infected at an ROI of 1 on 5DIV and analyzed on 11DIV. For *in vivo* applications, viral solutions were diluted to 1.0 × 10^8 viral particles/mL prior to stereotaxic delivery.

#### Stereotaxic injections

Sterotaxic injections followed previously described protocols (*54, 59*). Mice were placed in stereotaxic frames and anesthetized with 2% isoflurane (Patterson Veterinary; 2L per min oxygen flow rate). Fur near the incision was trimmed and ophthalmic eye ointment was applied to prevent desiccation for the cornea during surgery. Viral solutions (Gnaq(Q209L) or GFP control) were injected bilaterally into hippocampal CA1 and dentate gyrus using the following coordinates: (from bregma) anterior: -2 mm and lateral: 1.5 mm and (from skull surface) height: -1.7 mm and - 2.1 mm. A volume of 2 μL of viral solution was injected into each hemisphere over 10 min (injection speed of 0.1 μL per min) using a 5 μL 30s-guage Hamilton syringe. The needle was maintained in place for 8 min, slowly retracted half-way and kept in position for 2 min, and then removed, to prevent backflow along the injection tract. The incision was closed with a silk suture. Mice then received a subcutaneous injection of saline, enrofloxacin (Bayer) antibiotic, carprofen (Patterson) and buprenorphine (Butler Schein) for pain. Mice were housed alone and monitored during recovery. 14-16 mice were injected for each group.

#### Mouse behavior

##### Nestlet

The nestlet assay was completed to assess well-being following protocols described previously (*26*). Mice were provided with pressed cotton nestlets and given 48 hours to build nests. After 48 hours, a nestlet score (1-5) was given based on the quality of nests. A score of 1 = no nest built and a score of 5 = an enclosed nest (**Fig. 3F**). The scores from two separate nesting experiments performed at a one-week interval were averaged for each mouse.

##### Novel object recognition

The novel object recognition task was performed following the protocol previously described(*54, 59*). During habituation (day 1) mice could freely explore an empty, 40 cm x 40 cm square arena. During training (day 2), two identical objects were placed into the arena, and mice could explore the objects for 5 min. For testing (day 3), one object was replaced with a novel object, and mice could explore the objects for 5 min. Time spent exploring each object was quantified using Smart Video Tracking Software (Panlab; Harvard Apparatus). To control for any inherent object preference, two different sets of objects were used for experiments. To control for any potential object-independent location preference, the location of the novel object, relative to the trained object, was varied. Objects were selected based on their ability to capture mouse interest, regardless of genetic background or age. The time spent with the novel object was calculated by: ((Time with novel object)/(Time with trained object + Time with novel object)) * 100. In this object preference index, 100% indicates full preference for the novel object and 0% indicates full preference for the trained object. A score of 50% would mean the mouse spend equal times exploring both objects. If a mouse did not explore either object during training, they were excluded from the analysis.

##### Radial arm water maze

The radial arm water maze (RAWM) to assess spatial learning and memory was completed following previously described protocols (Alamed et al., 2006). For this task, the goal arm containing the platform remained in the same location for training and testing phases, but the starting arm was changed during each trial. Spatial cues were posted on all four walls in the RAWM area. Entry into the incorrect arm was scored as an error, and errors were averaged over training blocks (three consecutive trials). During the training phase (day 1), mice trained for 15 trials, with trials alternating between a visible and hidden platform for blocks 1-4 and then switching to only a hidden platform in block 5. During the testing phase (day 2) mice were tested for 15 trials with a hidden platform for blocks 6-10. Scoring was performed blind to treatment condition.

#### Tissue collection

Following the completion of behavioral assays, all mice were anesthetized with 87.5 mg/kg ketamine and 12.5mg/kg xylazine, transcardially perfused with ice-cold phosphate buffered saline and brains dissected. For one hemisphere, hippocampi were subdissected, flash frozen, and stored at - 80°C for snRNAseq analysis. The second hemisphere was post-fixed in 4% para-formaldehyde, pH 7.4, at 4°C for 48 hours before cryprotection with 30% sucrose. Brains were sectioned coronally at 40 μm with a freeze-stage microtome (Leica Camera, Inc.) and stored at -20°C in cryo-protective media until immunofluorescence, immunohistochemistry, or RNA-scope.

#### Immunofluorescence and microscopy

Immunofluorescence methods followed those described previously (*6, 54*). One hemisphere of each mouse brain was sectioned at 40 μm though the entire hippocampus and stored in cryoprotective media at -20°C until use. Sections for each mouse were stored in tubes, with each tube containing sections representative of the entire hippocampus. From each tube, 3-4 sections containing the dorsal hippocampus was selected for each immunofluorescence experiment. Sections were first washed 3x in TBST (Millipore Sigma, Cat. # 91414) for 5 min each, and then permeabilized with 0.1% Triton X-100 (Millipore Sigma, Cat. # 11332481001) for 30 min at room temperature. Sections were then washed 3x, for 5 min each in TBST and blocked in donkey serum (Southern Biotech, Cat. # 0030-01) for 1 hour at room temperature. Sections incubated shaking, overnight at 4°C in primary antibody solutions in TBST with 3% donkey serum. The dilutions were as follows: synapsin: 1:1000 (Anti-Synapsin I antibody; Abcam, Cat. # ab6458)1, PSD95: 1:500 (Anti-PSD95 antibody; Sigma-Aldrich, Cat. # P246), gephyrin: 1:200 (Gephryin antibody; Synaptic Systems, Cat. # 147 318), Arc: 1:500 (Arc (C-7); Santa Cruz Biotech, Cat. # sc-17839), and Erg1: 1:500 (Erg1 Antibody; Thermo Scientific, Cat. # MA5-15009). The next day, sections were washed 3x for 5 min each in TBST and then incubated in secondary antibody solution in TBST with 3% donkey serum, shaking for 1 hour at room temperature. All secondary antibodies were used at a 1:500 dilution and included the following: Anti-Mouse IgG (H+L), highly cross-adsorbed, CF 647 antibody (Sigma-Aldrich, Cat. # SAB4600176), Anti-Rabbit IgG (H+L), highly cross-adsorbed, CF 555 antibody (Sigma-Aldrich, SAB4600061), Alexa Fluor 647-AffiniPure Donkey Anti-Guinea Pig IgG (H+L) (Jackson ImmunoResearch, Cat. #706-605-148). All samples were then washed 3x in TBST and stained with Hoechst (1:1000 in TBST; Molecular Probes Hoechst 33342, Thermo Fisher, Cat. # H3570) for 10 min at room temperature. Samples were lastly washed 3x in TBST and mounted using Prolong Gold Medium (Molecular Probes, Thermo Fisher, Cat. #P36934). For each analysis, *n =* 12 mice per group.

For the synaptic proteins (protocol modified from (*60*)), slides were imaged using a Nikon A1R Confocal Microscope equipped with a 60x oil objective. Z-stacks through the entire section, spaced at 0.33 μm apart, were taken from the same region of DG and CA1 from all hippocampal sections for each mouse on 3-4 different sections. Using FIJI, Z-stacks were analyzed by max intensity projections for clusters of 3 consecutive stacks, to avoid false positives. A macro was written to identify colocalized synaptic puncta with the following processing parameters: despeckle, subtract background (rolling ball, radius of 10 pixels), thresholding with triangle method, and watershed to separate overlapping particles. Colocalized and non-colocalized points were then counted using the colocalization plugin followed by analyze particles. An average for synapsin, gephyrin, PSD95, and excitatory and inhibitory synapses was calculated for each mouse by averaging values across sections.

To image immediate early genes Arc and Erg1, a Spinning Disk CSU-W1 Nikon Confocal Microscope (Princeton University Microscopy Core) with a 20x objective was used, and z-stacks spaced 3 μm apart were taken through the entire section and stitched together to include the entire hippocampus. After max intensity projection, FIJI was used for analysis. The ROI (CA1 or DG) was outlined in each section. To identify the percent positive cells per ROI the following parameters were used: Gaussian filtering (sigma of 2), thresholding using the triangle method, and counting particles greater than 15 pixels. This value was then divided by the ROI area sampled. Mean signal (normalized to Hoechst) was measured when there were not punctate particles to count. For all analyses, an average was calculated for each mouse across all sections for CA1 and DG.

#### Neuronal nuclei isolation

Neuronal nuclei were isolated similar to previously described methods (*54, 61*). One hemisphere of the hippocampus dissected, flash frozen, and stored at -80°C until use. Samples were kept on ice during the experiment (*n =* 4 GFP hippocampi and 4 GNAQ hippocampi). The hippocampi were dounce homogenized using 1 mL dounce tissue grinders (Wheaton, DWK Life Sciences, Cat. # 357538) in 300 L of NP40 lysis buffer (10 mM Tris-HCl (Sigma-Aldrich, Cat. # T2194; pH 7.4), 10 mM NaCl (Sigma-Aldrich, Cat. # 59222C; 5M), 3 mM MgCl_2_ (Sigma-Aldrich, Cat. # M1028; 1M), 0.1% Nonidet P40 Substitute (Sigma-Aldrich, Cat. # 74385), 1 mM DTT (Sigma-Aldrich, Cat. # 646563), 1 U/μL RNase inhibitor (Sigma Protector RNase inhibitor; Sigma-Aldrich, Cat. # 3335402001), Nuclease-free water) with 15 loose stokes followed by 15 tight strokes. Samples were incubated for 7 min on ice, and pipetted twice after 2 min and 5 min, 3x at each point, with a P200 pipette. Suspensions were passed through a 40 μm filter into a 15 mL conical tube, and then transferred to a 2-mL low bind microcentrifuge tube. Samples were centrifuged at 500 rcf for 5 min at 4°C. The supernatant was removed and 1 mL of wash buffer (PBS + 1% BSA (Miltenyi Biotec, Cat. # 130-091-376) + 1 U/uL of RNAase inhibitor) was added with no mixing for 5 min on ice. The pellet was then resuspended by pipette mixing and samples were centrifuged again at 500 rcf for 5 min at 4°C. The supernatant was removed and the cells were resuspended in 1 mL of the same wash buffer. Next, 10 μL of Hoechst stain (1:10,000 dilution; Molecular Probes Hoechst 33342, Thermo Fisher, Cat. # H3570) was added to each sample, prior to filtering into the FACS tube and incubating on ice for 5 min on the way to the FACS core.

#### FACS, snRNA-seq library preparation, and sequencing

The Princeton FACS Core helped sort Hoechst and GFP+ nuclei using a 70 μm nozzle and a flow rate of 3 on a BD Biosciences FACSAria Fusion sorter into a 1.5 mL low bind microcentrifuge tube containing collection buffer (500 μL of 1% BSA + 1.5 U/μL RNase Inhibitor).

After FACS, samples were centrifuged at 500 rcf for 5 min at 4°C. The supernatant was removed and nuclei were resuspended in 20 μL of collection buffer. For each sample, 2 μL of nuclei were diluted 1:5 in PBS + Trypan Blue (Gibco Trypan Blue Solution, Fisher Scientific, Cat. # 15-250-061) and counted on a hemocytometer (Millicell Disposable Hemocytometer, Millipore Sigma, Cat. # MDH-2N1-50PK). The number of nuclei/μL was estimated for each sample and provided to the Princeton Genomics Core. Single nuclei suspension samples were loaded to the 10X Genomics Chromium X system using the Single Cell 3’ v3.1 Reagent Kits (10X Genomics Inc., CA) to generate and amplify cDNA. The amplified cDNA samples were purified with Ampure XP magnetic beads (Beckman Coulter, CA), quantified by Qubit fluorometer (Invitrogen, CA), and examined on Bioanalyzer with High Sensitivity DNA chips (Agilent, CA) for size distribution. Illumina sequencing libraries were generated from the amplified cDNA samples using the Illumina Tagment DNA Enzyme and Buffer kit (Illumina, CA). These libraries were examined by Qubit and Bioanalyzer, then pooled at equal molar amount and sequenced on Illumina NovaSeq 6000 S Prime flowcells as 28+94 nt pair-end reads following the standard protocol. Raw sequencing reads were filtered by Illumina NovaSeq Control Software and only the Pass-Filter (PF) reads were used for further analysis.

#### snRNA-seq analysis

##### Raw Data Handling

After sequencing, output fastq files were mapped to an mm10 reference genome that was edited to include eGFP and filtered using Cell Ranger version 7.1.0. The pooled samples consisted of GFP control samples: four samples, pooled in groups of two, and GNAQ samples: four samples, pooled in groups of two. For the GFP control samples, Cell Ranger estimated 12,771 cells or 8,852 cells for the two pooled samples, with ∼4500 median UMIs per cell. In the GNAQ samples, Cell Ranger estimated 10,860 and 9,832 cells in each of the two pooled samples, with ∼4000 median UMIs per cell.

##### Quality Control

Ambient RNA contamination was detected and removed using the SoupX package in R. Full Cell Ranger out files were inputted and ambient mRNA expression profile was estimated from empty droplets, the contamination fraction of each cell was calculated, and then the UMIs for all cells were corrected based on the contamination fraction. The global contamination fraction of all samples was estimated to be 0.03% which falls between the normal range of 0.02-0.05%. Note: SoupX removes ambient RNA contamination, it does not remove any cells from the dataset.

Additional quality control was done in R using the Seurat package. Violin Plots of the nFeatures/cell and the mtRNA/cell were generated, and cells were removed based on Seurat’s suggested cutoffs. For our samples, cells with <200 nFeatures (potentially damaged cells) or >5000 nFeatures (potential doublets) were filtered out. Cells with mitochondrial contamination >0.25% were also removed from the dataset.

Following quality control, our final cell numbers for GFP control samples were 12,227 and 8,183. Our two GNAQ samples had final cell numbers of 9,325 and 10,231 cells. Approximately 500-600 cells were filtered out of each condition after all quality control.

##### Normalization, Integration, and Clustering

Prior to normalization, all datasets were merged into a single Seurat object. The data was normalized using SCTransform to fit the gene expression counts along a negative binomial distribution. 3000 integration features were selected from the dataset and used to find integration anchors. Integration was then performed on the SCTransform normalized counts.

Next, principal component analysis was performed, and an elbow plot of the principal components was generated to help choose the number of components needed to accurately represent the data. We chose 40 principal components to perform clustering. Clustering was performed using the Louvain algorithm for network clustering at a resolution of 0.5 which resulted in 31 clusters.

##### Cell Type Annotation

Finally, clusters were annotated using known hippocampal markers for different cell types (Allen Mouse Brain Atlas (http://mouse.brain-map.org/), HippoSeq (https://hipposeq.janelia.org/), (*62*–*64*)). The markers are in **Table S2**.

#### Data and statistical analyses

For worm experiments, data are expressed at mean ± s.e.m. Statistical analysis was performed with Prism 8.0 or 9.0 software (GraphPad Software). Means between two groups were compared with two-tailed, unpaired Student’s *t*-test. Comparisons of means from multiple groups with each other were analyzed with one-way ANOVA followed by a Bonferroni *post hoc* test, as indicated in the figure legends. Additional statistical details are reported in figure legends.

All mouse experiments were randomized and blinded by an independent researcher before behavioral testing. Researchers remained blinded throughout histological, biochemical and behavioral assessments. Groups were un-blinded at the end of each experiment upon statistical analysis. Data are expressed as mean ± s.e.m. The distribution of data in each set of experiments was tested for normality using D’Agostino-Pearson omnibus test or Shapiro-Wilk test. Statistical analysis was performed with Prism 8.0 or 9.0 software (GraphPad Software). Means between two groups were compared with two-tailed, unpaired Student’s *t*-test. Comparisons of means from multiple groups with each other were analyzed with one-way ANOVA followed by the appropriate *post hoc* test, as indicated in the figure legends. Trial by group interactions were analyzed using repeated measures ANOVA and with Šídák’s correction for multiple comparisons. Additional statistic details are indicated in the respective figure legends. All data generated or analyzed in this study are included in this article.

## Supplementary Tables and Figures

**Table S1: Differential expression master list**. List of all cell types and significantly upregulated genes with GNAQ treatment, relative to GFP control. Mouse gene names, gene descriptions, and worm orthologs are also listed.

**Table S2: Genes used to identify clusters**. List of genes that were primarily used to define clusters based on (ref. 57-59).

**Fig. S1.**
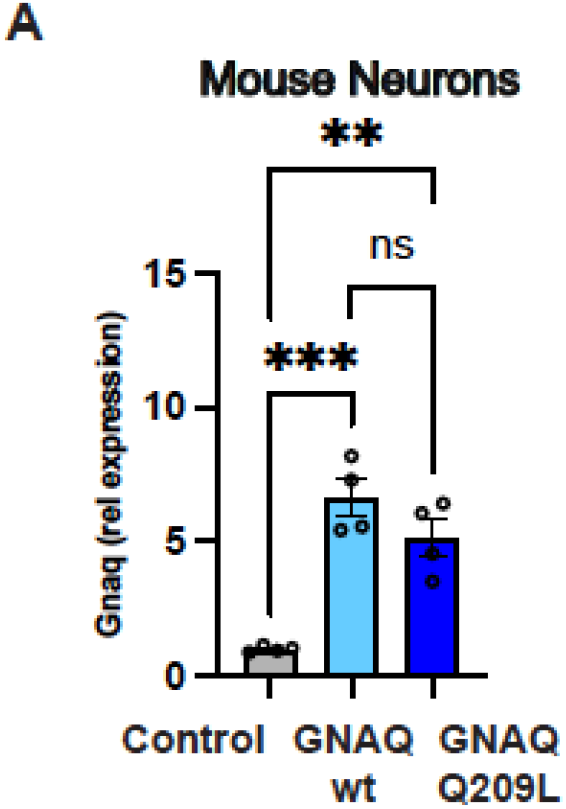
Relative Expression of *Gnaq* in mouse primary neurons. **(A)** *Gnaq* expression for GNAQ wt and GNAQ(Q209L) were elevated compared to control, but they did not differ from each other.

## Notes

### Competing Interest Statement

The authors have declared no competing interest.

